# The *Clostridioides difficile* species problem: global phylogenomic analysis uncovers three ancient, toxigenic, genomospecies

**DOI:** 10.1101/2020.09.21.307223

**Authors:** Daniel R. Knight, Korakrit Imwattana, Brian Kullin, Enzo Guerrero-Araya, Daniel Paredes-Sabja, Xavier Didelot, Kate E. Dingle, David W. Eyre, César Rodríguez, Thomas V. Riley

## Abstract

*Clostridioides difficile* infection (CDI) remains an urgent global One Health threat. The genetic heterogeneity seen across *C. difficile* underscores its wide ecological versatility and has driven the significant changes in CDI epidemiology seen in the last 20 years. We analysed an international collection of over 12,000 *C. difficile* genomes spanning the eight currently defined phylogenetic clades. Through whole-genome average nucleotide identity, pangenomic and Bayesian analyses, we identified major taxonomic incoherence with clear species boundaries for each of the recently described cryptic clades CI-III. The emergence of these three novel genomospecies predates clades C1-5 by millions of years, rewriting the global population structure of *C. difficile* specifically and taxonomy of the *Peptostreptococcaceae* in general. These genomospecies all show unique and highly divergent toxin gene architecture, advancing our understanding of the evolution of *C. difficile* and close relatives. Beyond the taxonomic ramifications, this work impacts the diagnosis of CDI worldwide.

## Introduction

The bacterial species concept remains controversial, yet it serves as a critical framework for all aspects of modern microbiology^1^. The prevailing species definition describes a genomically coherent group of strains sharing high similarity in many independent phenotypic and ecological properties^2^. The era of whole-genome sequencing (WGS) has seen average nucleotide identity (ANI) replace DNA-DNA hybridization as the ‘next-generation’ standard for microbial taxonomy^3, 4^. Endorsed by the National Center for Biotechnology Information (NCBI)^4^, ANI provides a precise, objective and scalable method for delineation of species, defined as monophyletic groups of strains with genomes that exhibit at least 96% ANI^5, 6^.

*Clostridioides (Clostridium) difficile* is an important gastrointestinal pathogen that places a significant growing burden on health care systems in many regions of the world^7^. In both its 2013^8^ and 2019^9^ reports on antimicrobial resistance (AMR), the US Centers for Disease Control and Prevention rated *C. difficile* infection (CDI) as an urgent health threat, the highest level. Community-associated CDI has become more frequent^7^, likely because *C. difficile* has become established in livestock worldwide, resulting in significant environmental contamination^10^. Thus, over the last two decades, CDI has emerged as an important One Health issue^10^.

Based on multi-locus sequence type (MLST), there are eight recognised monophyletic groups or ‘clades’ of *C. difficile*^11^. Strains within these clades show many unique clinical, microbiological and ecological features^11^. Critical to the pathogenesis of CDI is the expression of the large clostridial toxins, TcdA and TcdB and, in some strains, binary toxin (CDT), encoded by two separate chromosomal loci, the PaLoc and CdtLoc, respectively^12^. Clade 1 (C1) contains over 200 toxigenic and non-toxigenic sequence types (STs) including many of the most prevalent strains causing CDI worldwide e.g. ST2, ST8, and ST17^11^. Several highly virulent CDT-producing strains, including ST1 (PCR ribotype (RT) 027), a lineage associated with major hospital outbreaks in North America, Europe and Latin America^13^, are found in clade 2 (C2). Comparatively little is known about clade 3 (C3) although it contains ST5 (RT 023), a toxigenic CDT-producing strain with characteristics that may make laboratory detection difficult^14^. *C. difficile* ST37 (RT 017) is found in clade 4 (C4) and, despite the absence of a toxin A gene, is responsible for much of the endemic CDI burden in Asia^15^. Clade 5 (C5) contains several CDT-producing strains including ST11 (RTs 078, 126 and others), which are highly prevalent in production animals worldwide^16^. The remaining so-called ‘cryptic’ clades (C-I, C-II and C-III), first described in 2012^17, 18^, contain over 50 STs from clinical and environmental sources^17, 18, 19, 20, 21^. Evolution of the cryptic clades is poorly understood. Clade C-I strains can cause CDI, however, due to atypical toxin gene architecture, they may not be detected, thus their prevalence may have been underestimated^21^.

There are over 600 STs currently described and some STs may have access to a gene pool in excess of 10,000 genes^11, 16, 16^. Considering such enormous diversity, and recent contentious taxonomic revisions^23, 24^, we hypothesise that *C. difficile* comprises a complex of distinct species divided along the major evolutionary clades. In this study, whole-genome ANI, and pangenomic and Bayesian analyses are used to explore an international collection of over 12,000 *C. difficile* genomes, to provide new insights into ancestry, genetic diversity and evolution of pathogenicity in this enigmatic pathogen.

## Results

### An updated global population structure based on sequence typing of 12,000 genomes

We obtained and determined the ST and clade for a collection of 12,621 *C. difficile* genomes (taxid ID 1496, Illumina data) existing in the NCBI Sequence Read Archive (SRA) as of 1^st^ January 2020. A total of 272 STs were identified spanning the eight currently described clades, indicating that the SRA contains genomes for almost 40% of known *C. difficile* STs worldwide (n=659, PubMLST, January 2020). C1 STs dominated the database in both prevalence and diversity (**Fig. 1**) with 149 C1 STs comprising 57.2% of genomes, followed by C2 (35 STs, 22.9%), C5 (18 STs, 10.2%), C4 (34 STs, 7.5%), C3 (7 STs, 2.0%) and the cryptic clades C-I, C-II and C-III (collectively 17 STs, 0.2%). The five most prevalent STs represented were ST1 (20.9% of genomes), ST11 (9.8%), ST2 (9.5%), ST37 (6.5%) and ST8 (5.2%), all prominent lineages associated with CDI worldwide^11^.

**Figure 1.**
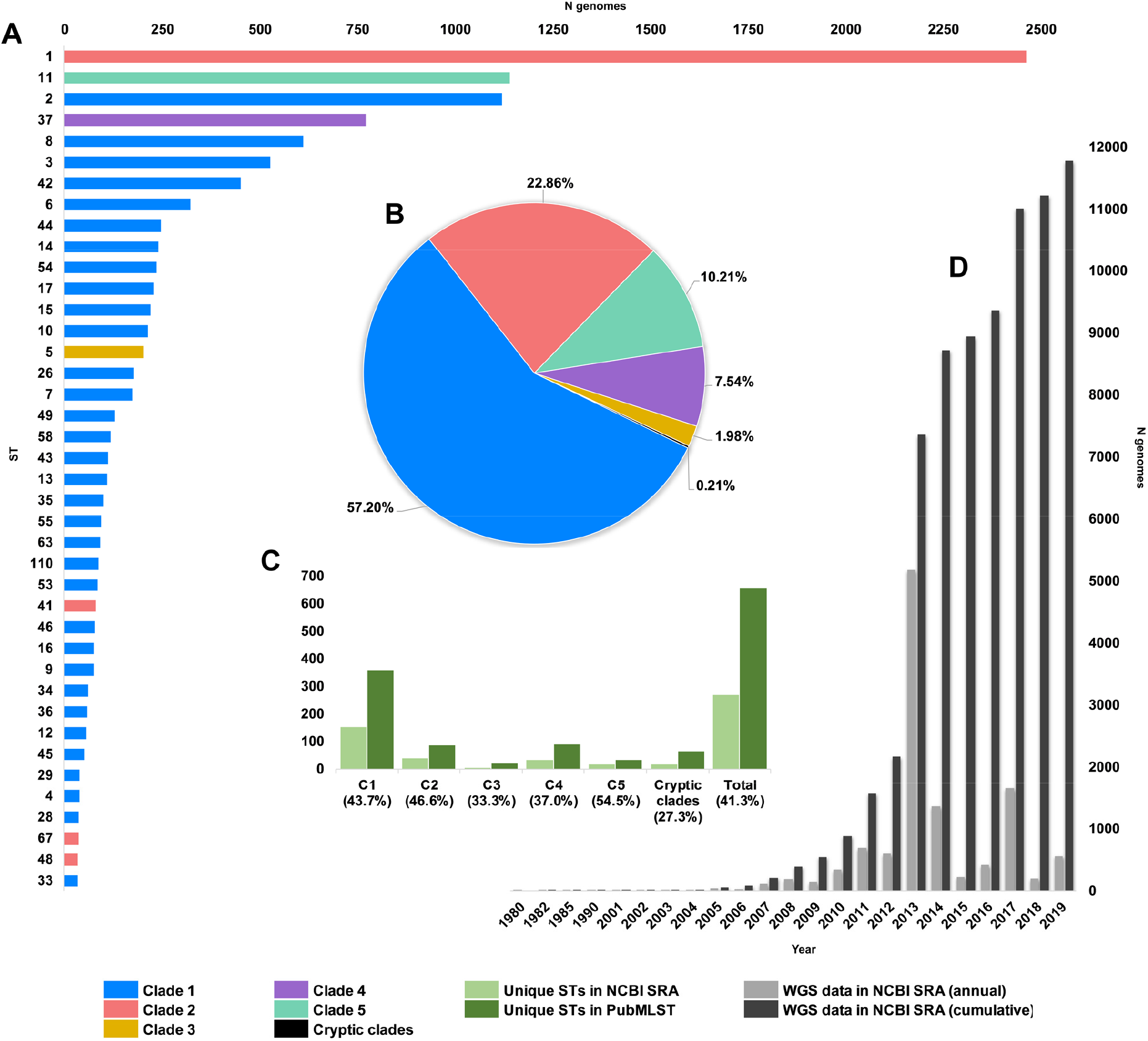
Composition of *C. difficile* genomes in the NCBI SRA. Snapshot obtained 1^st^ January 2020; 12,304 strains, [taxid ID 1496]. **(A)** Top 40 most prevalent STs in the NCBI SRA coloured by clade. **(B)** The proportion of genomes in ENA by clade. **(C)** Number/ proportion of STs per clade found in the SRA/present in the PubMLST database. **(D)** Annual and cumulative deposition of *C. difficile* genome data in ENA.

**Fig. 2** shows an updated global *C. difficile* population structure based on the 659 STs; 27 novel STs were found (an increase of 4%) and some corrections to assignments within C1 and C2 were made, including assigning ST122^25^ to C1. Based on PubMLST data and bootstraps values of 1.0 in all monophyletic nodes of the cryptic clades (**Fig. 2**), we could confidently assign 25, 9 and 10 STs to cryptic clades I, II and III, respectively. There remained 26 STs spread across the phylogeny that did not fit within a specific clade (defined as outliers). The tree file for **Fig. 2** and full MLST data is available as **Supplementary Data** at http://doi.org/10.6084/m9.figshare.12471461.

**Figure 2.**
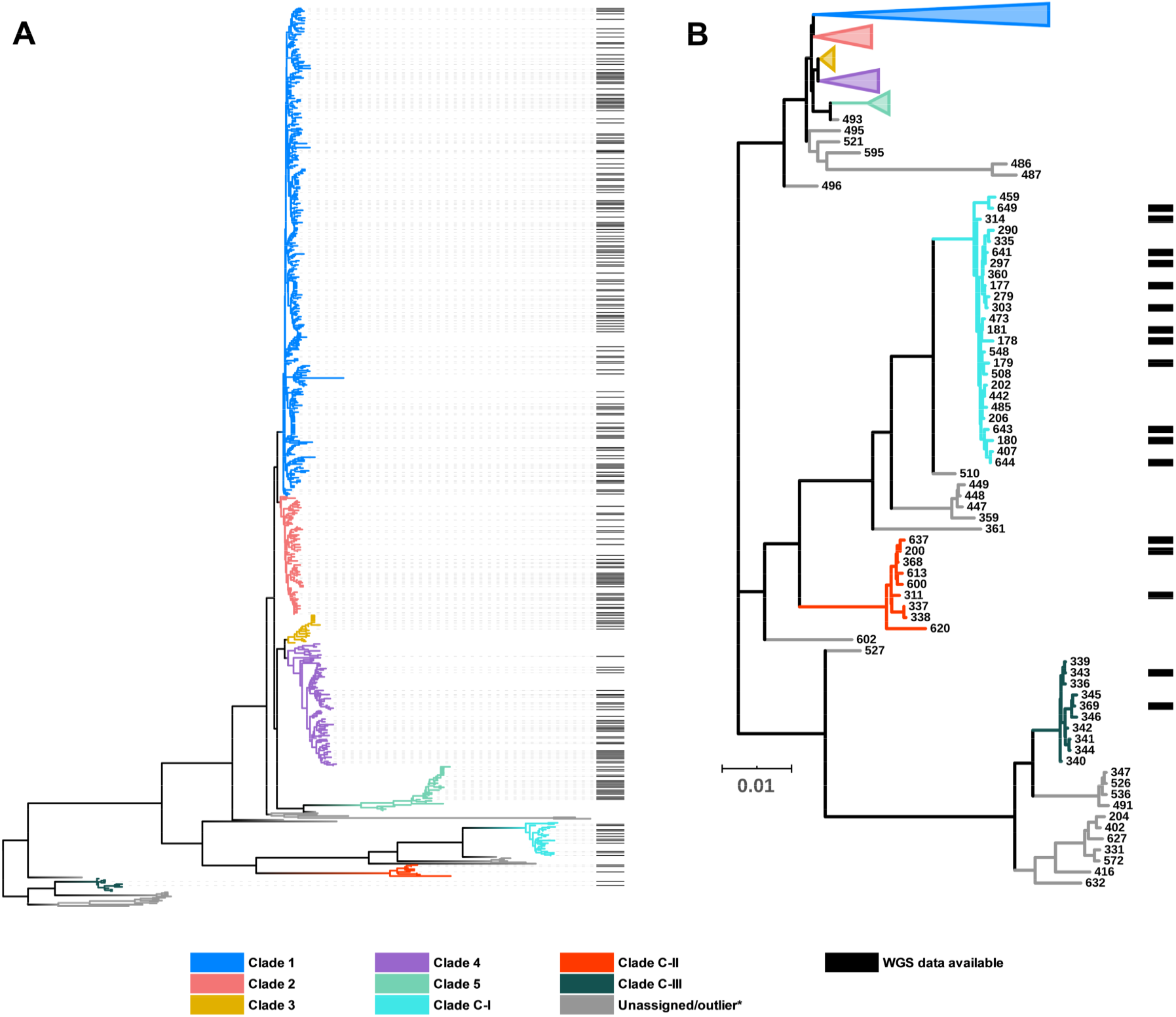
*C. difficile* population structure. **(A)** NJ phylogeny of 659 aligned, concatenated, multilocus sequence type allele combinations coloured by current PubMLST clade assignment. Black bars indicate WGS available for ANI analysis (n=260). **(B)** A subset of the NJ tree showing cryptic clades C-I, C-II and C-III. Again, black bars indicate WGS available for ANI analysis (n=17).

### Whole-genome ANI analysis reveals clear species boundaries

Whole-genome ANI analyses were used to investigate genetic discontinuity across the *C. difficile* species (**Fig. 3** and **Supplementary Data**). Representative genomes of each ST, chosen based on metadata, read depth and quality, were assembled and annotated. Whole-genome ANI values were determined for a final set of 260 STs using three independent ANI algorithms (FastANI, ANIm and ANIb, see *Methods*). All 225 genomes belonging to clades C1-4 clustered within an ANI range of 97.1-99.8% (median FastANI values of 99.2, 98.7, 97.9 and 97.8%, respectively, **Fig. 3A-C**).

**Figure 3.**
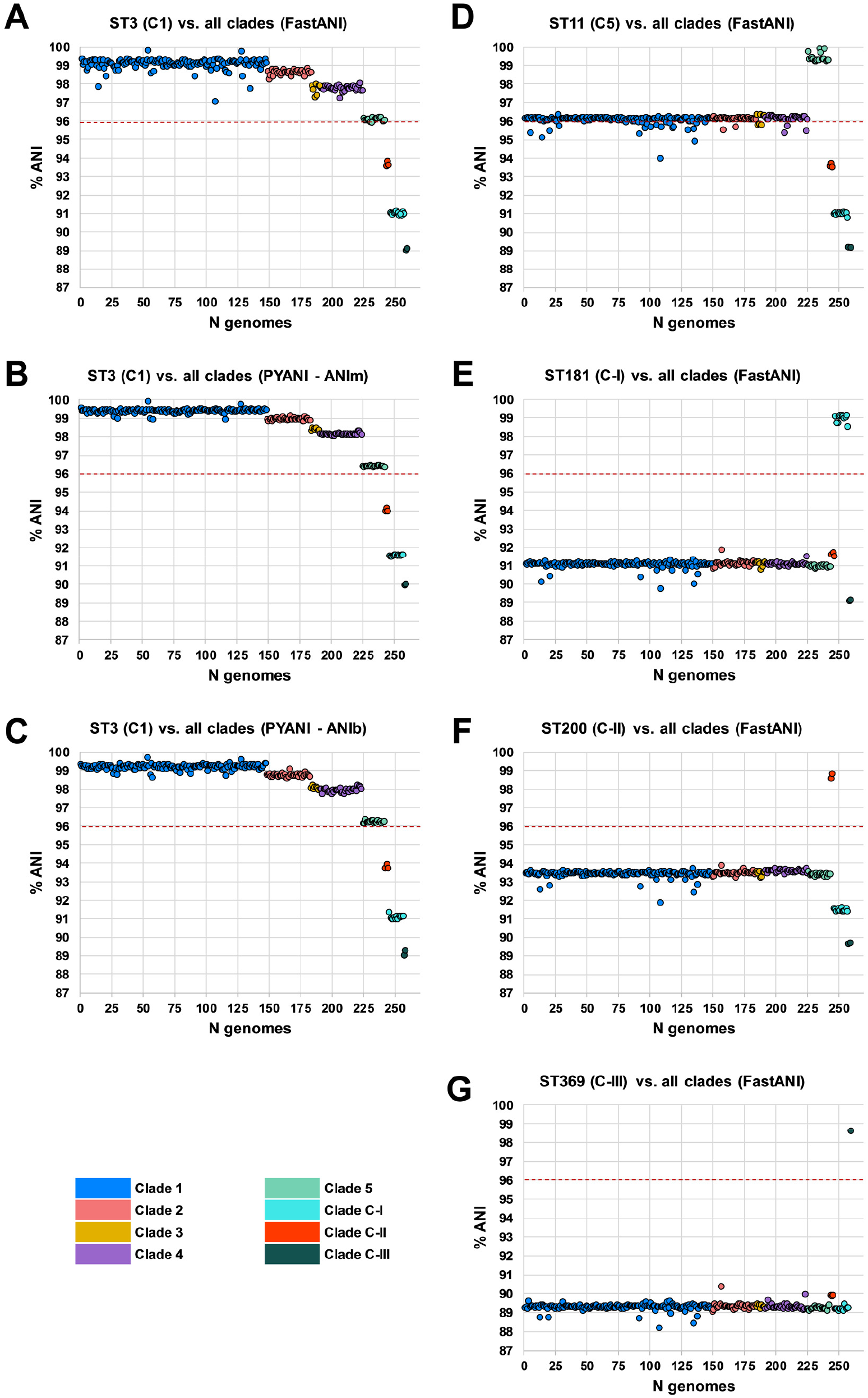
Species-wide ANI analysis. Panels **A-C** show ANI plots for ST3 (C1) vs. all clades (260 STs) using FastANI, ANIm and ANIb algorithms, respectively. Panels **D**-**G** show ANI plots for ST11 (C5), ST181 (C-I), ST200 (C-II) and ST369 (C-III) vs all clades (260 STs), respectively. NCBI species demarcation of 96% indicated by red dashed line^4^.

These ANI values are above the 96% species demarcation threshold used by the NCBI^4^ and indicate that strains from these clades belong to the same species. ANI values for all 18 genomes belonging to C5 clustered on the borderline of the species demarcation threshold (FastANI range 95.9-96.2%, median 96.1%). ANI values for all three cryptic clades fell well below the species threshold; C-I (FastANI range 90.9-91.1%, median 91.0%), C-II (FastANI range 93.6-93.9%, median 93.7%) and C-III (FastANI range 89.1-89.1%, median 89.1%). All results were corroborated across the three independent ANI algorithms (**Fig. 3A-C**). *C. difficile* strain ATCC 9689 (ST3, C1) was defined by Lawson *et al*. as the type strain for the species^23^, and used as a reference in all the above analyses. To better understand the diversity among the divergent clades themselves, FastANI analyses were repeated using STs 11, 181, 200 and 369 as reference archetypes of clades C5, C-I, C-II and C-III, respectively. This approach confirmed that C5 and the three cryptic clades were as distinct from each other as they were collectively from C1-4 (**Fig. 3D-G**).

### Taxonomic placement of cryptic clades predates *C. difficile* emergence by millions of years

Previous studies using BEAST have estimated the common ancestor of C1-5 existed between 1 to 85 or 12 to 14 million years ago (mya)^26, 27^. Here, we used an alternative Bayesian approach, BactDating, to estimate the age of all eight *C. difficile* clades currently described. The last common ancestor for *C. difficile* clades C1-5 was estimated to have existed ∼3.89 mya with a 95% credible interval (CI) of 1.11 to 6.71 mya (**Fig. 4**). In contrast, C-II, C-I and C-III emerged 13.05 mya (95% CI 3.72-22.44), 22.02 (95% CI 6.28-37.83) and 47.61 mya (95% CI 13.58-81.73), respectively, at least 9 million years (Megaannum, Ma) before the common ancestor of C1-5. Independent analysis with BEAST, using a smaller core gene dataset (see *Methods*), provided broader estimates of clade emergence, though the emergence order was maintained; C1-5 12.01 mya (95% CI 6.80-33.47), C-II 37.12 mya (95% CI 20.95-103.48), C-I 65.93 mya (95% CI 37.32-183.84) and C-III 142.13 mya (95% CI 79.77-397.18) (**Fig. 4**).

**Figure 4.**
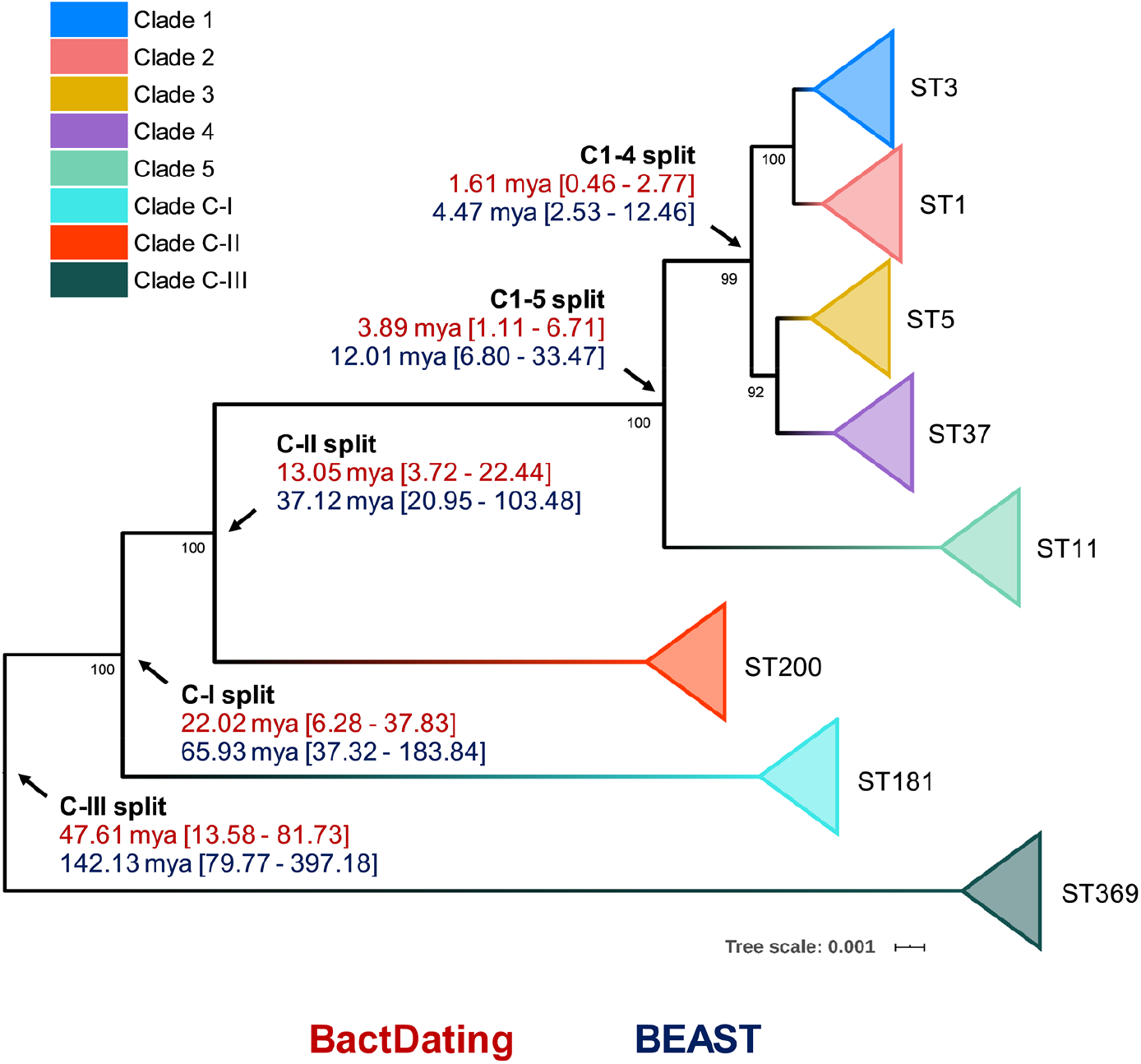
Bayesian analysis of species and clade divergence. BactDating and BEAST estimates of the age of major *C. difficile* clades. Node dating ranges for both Bayesian approaches are transposed onto an ML phylogeny built from concatenated MLST alleles of a dozen STs from each clade. Archetypal STs in each evolutionary clade are indicated. The tree is midpoint rooted and bootstrap values are shown. Scale bar indicates the number of substitutions per site. BactDating places the time of most recent common ancestor of C1-5 at 3.89 million years ago (mya) [95% credible interval (CI), 1.11-6.71 mya]. Of the cryptic clades, C-II shared the most recent common ancestor with C1-5 13.05 mya [95% CI 3.72-22.44 mya], followed by C-I (22.02 mya [95% CI 6.28-37.83 mya]), and C-III (47.61 mya [95% CI 13.58-81.73 mya]). Comparative estimates from BEAST are clades C1-5 (12.01 mya [95% CI 6.80-33.47 mya]), C-II (37.12 mya [95% CI 20.95-103.48 mya]), C-I (65.93 mya [95% CI 37.32-183.84 mya]), and C-III (142.13 [95% CI 79.77-397.18 mya]).

Next, to identify their true taxonomic placement, ANI was determined for ST181 (C-I), ST200 (C-II) and ST369 (C-III) against two reference datasets. The first dataset comprised 25 species belonging to the *Peptostreptococcaceae* as defined by Lawson *et al*.^23^ in their 2016 reclassification of *Clostridium difficile* to *Clostridioides difficile*. The second dataset comprised 5,895 complete genomes across 21 phyla from the NCBI RefSeq database (accessed 14^th^ January 2020), including 1,366 genomes belonging to *Firmicutes*, 92 genomes belonging to 15 genera within the *Clostridiales* and 20 *Clostridium* and *Clostridioides* species. The nearest ANI matches to species within the *Peptostreptococcaceae* dataset were *C. difficile* (range 89.3-93.5% ANI), *Asaccharospora irregularis* (78.9-79.0% ANI) and *Romboutsia lituseburensis* (78.4-78.7% ANI). Notably, *Clostridioides mangenotii*, the only other known member of *Clostridioides*, shared only 77.2-77.8% ANI with the cryptic clade genomes (**Table 1**).

**Table 1.**
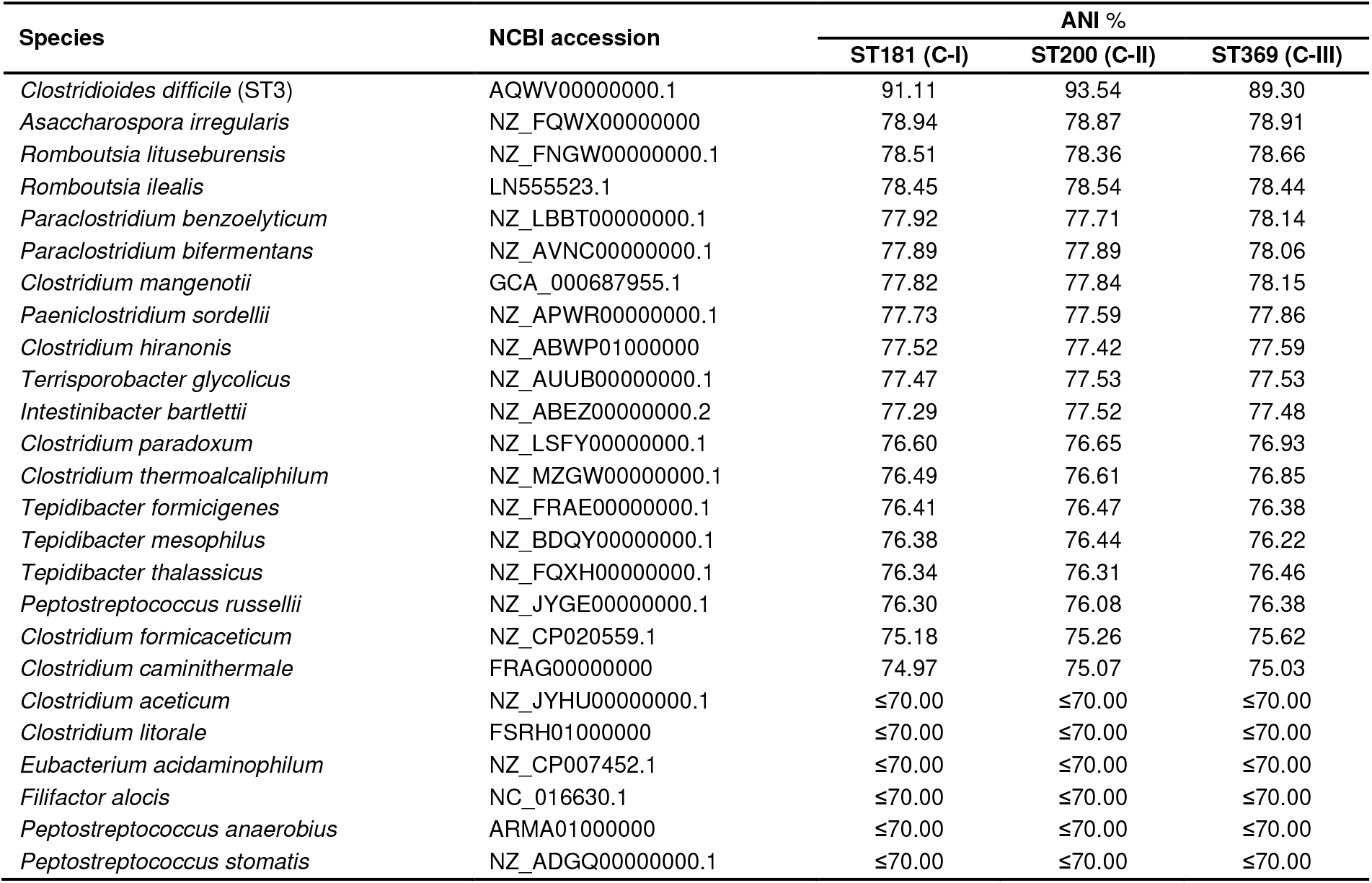
Whole-genome ANI analysis of cryptic clades vs. 25 *Peptostreptococcaceae* species from Lawson *et al*^23^.

Similarly, the nearest ANI matches to species within the RefSeq dataset were several *C. difficile* strains (range C-I: 90.9-91.1%; C-II: 93.4-93.6%; and C-III: 89.2-89.4%) and *Paeniclostridium sordellii* (77.7-77.9%). A low ANI (range ≤70-75%) was observed between the cryptic clade genomes and 20 members of the *Clostridium* including *C. tetani, C. botulinum, C. perfringens* and *C. butyricum*, the type strain of the *Clostridium* genus *senso stricto*. An updated ANI-based taxonomy for the *Peptostreptococcaceae* is shown in **Fig. 5A**. The phylogeny places C-I, C-II and C-III between *C. mangenotii* and *C. difficile* C1-5, suggesting that they should be assigned to the *Clostridioides* genus, distinct from both *C. mangenotii* and *C. difficile*. Comparative analysis of ANI and 16S rRNA values for the eight *C. difficile* clades and *C. mangenotii* shows significant incongruence between the data generated by the two approaches (**Fig. 5B**). The range of 16S rRNA % similarity between *C. difficile* C1-4, cryptic clades I-III and *C. mangenotii* was narrower (range 94.5-100) compared to the range of ANI values (range 77.8-98.7).

**Figure 5.**
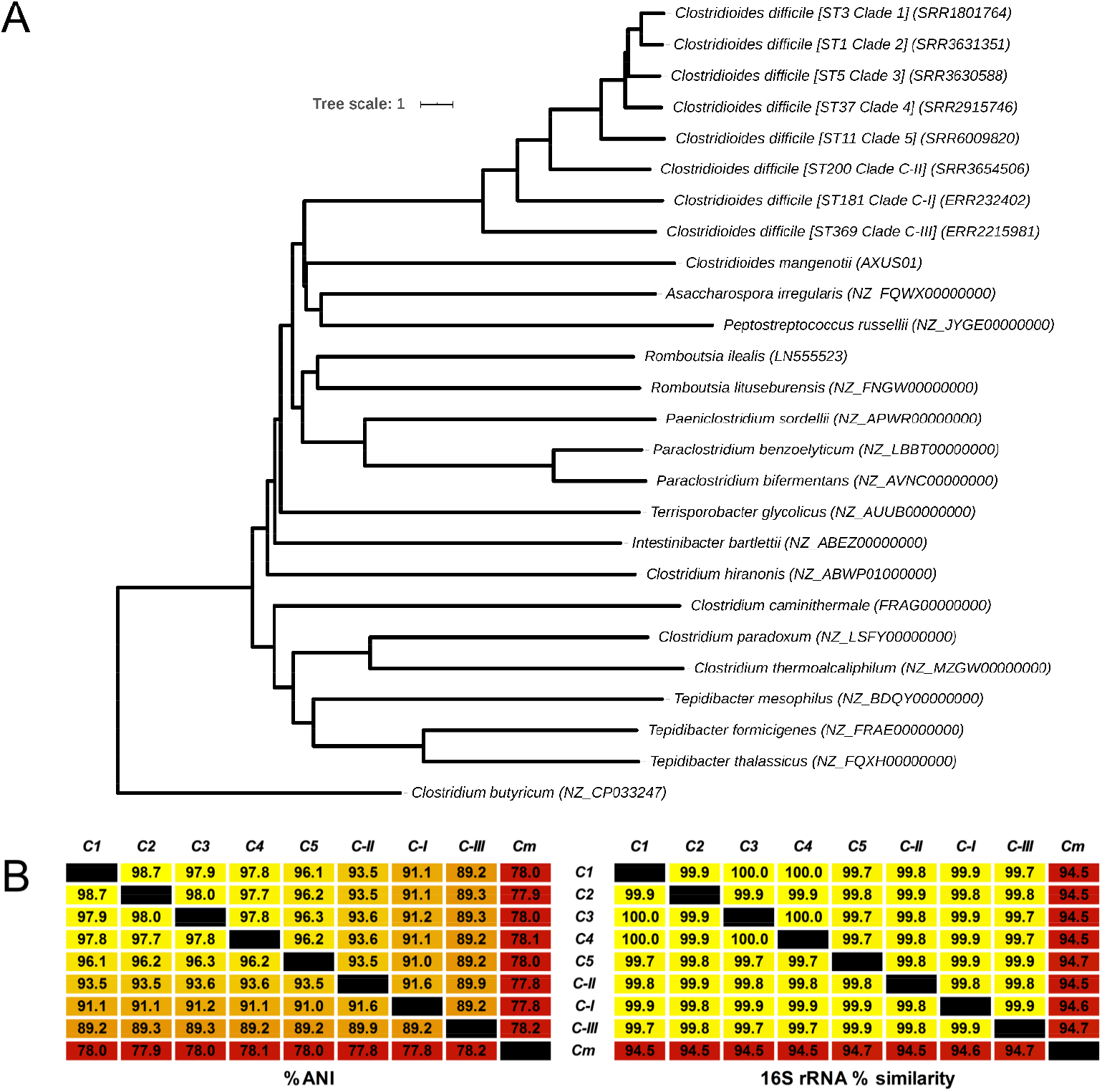
Revised taxonomy for the *Peptostreptococcaceae*. **(A)** ANI-based minimum evolution tree showing evolutionary relationship between eight *C. difficile* ‘clades’ along with 17 members of the *Peptostreptococcaceae* (from Lawson *et al*^23^) as well as *Clostridium butyricum* as the outgroup and type strain of the *Clostridium* genus *senso stricto*. To convert the ANI into a distance, its complement to 1 was taken. Matrices showing pairwise ANI and 16S rRNA values for the eight *C. difficile* clades and *C. mangenotii*, the only other known member of *Clostridioides*.

### Evolutionary and ecological insights from the *C. difficile* species pangenome

Next, we sought to quantify the *C. difficile* species pangenome and identify genetic loci that are significantly associated with the taxonomically divergent clades. With Panaroo, the *C. difficile* species pangenome comprised 17,470 genes, encompassing an accessory genome of 15,238 genes and a core genome of 2,232 genes, just 12.8% of the total gene repertoire (**Fig 6**). The size of the pangenome reduced by 2,082 genes with the exclusion of clades CI-III, and a further 519 genes with the exclusion of C5. Compared to Panaroo, Roary overestimated the size of the pangenome (32,802 genes), resulting in markedly different estimates of the percentage core genome, 3.9 and 12.8%, respectively (χ^2^=1,395.3, df=1, p<0.00001). Panaroo can account for errors introduced during assembly and annotation, thus polishing the 260 Prokka-annotated genomes with Panaroo resulted in a significant reduction in gene content per genome (median 2.48%; 92 genes, range 1.24-12.40%; 82-107 genes, p<0.00001). The *C. difficile* species pangenome was determined to be open^28^ (**Fig 6**).

**Figure 6.**
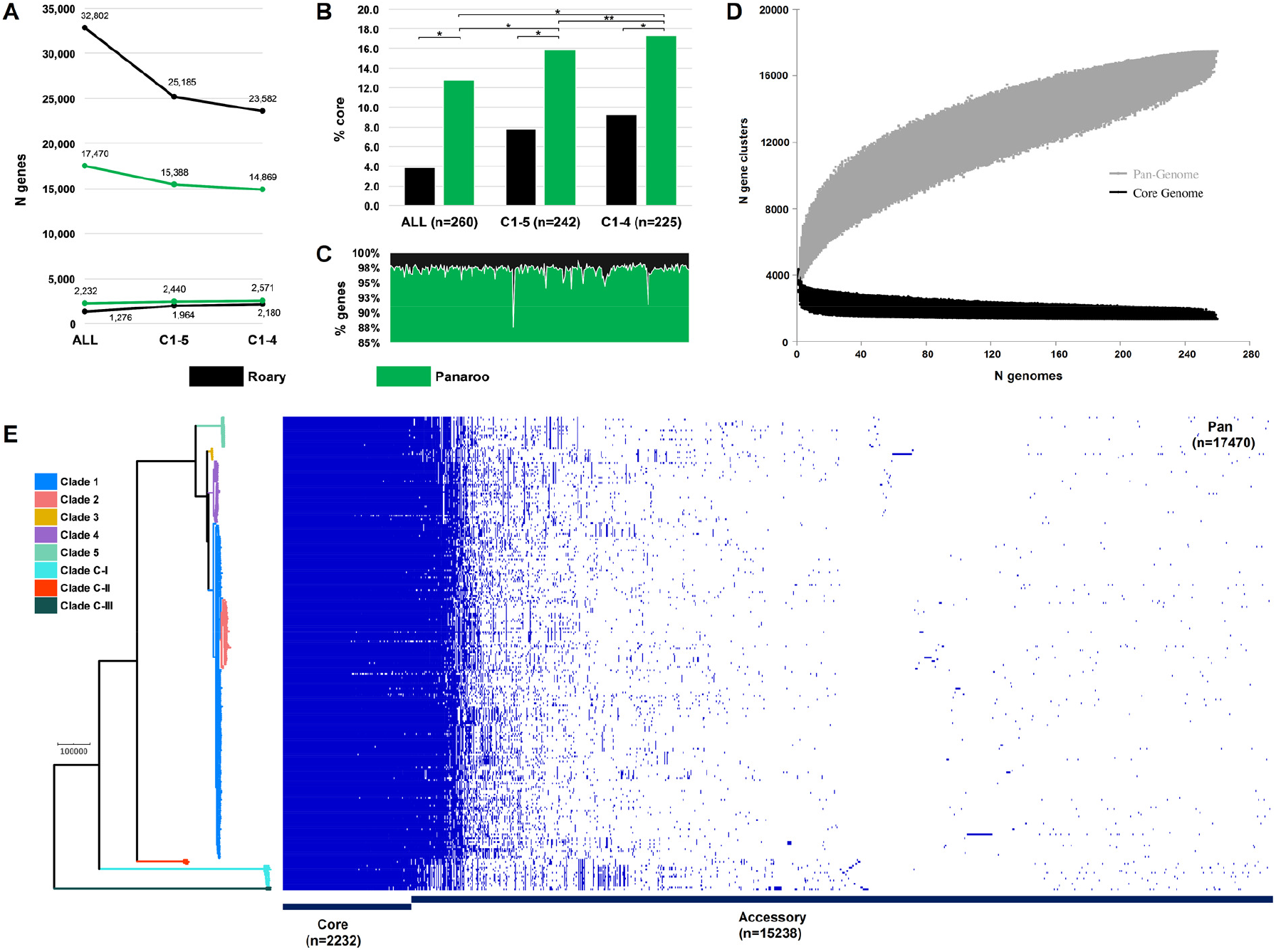
*Clostridioides difficile* species pangenome. **(A)** Pan and core genome estimates for all 260 STs, clades C1-4 (n=242 STs) and clades C1-5 (n=225 STs). **(B)** The difference in % core genome and pangenome sizes with Panaroo and Roary algorithms. (*) indicates χ^2^ p < 0.00001 and (**) indicates χ^2^ p = 0.0008. **(C)** The proportion of retained genes per genome after polishing Prokka-annotated genomes with Panaroo. **(D)** The total number of genes in the pan (grey) and core (black) genomes are plotted as a function of the number of genomes sequentially added (n=260). Following the definition of Tettelin *et al*.^28^, the *C. difficile* species pangenome showed characteristics of an “open” pangenome. First, the pangenome increased in size exponentially with sampling of new genomes. At n*=*260, the pangenome exceeded more than double the average number of genes found in a single *C. difficile* genome (∼3,700) and the curve was yet to reach a plateau or exponentially decay, indicating more sequenced strains are needed to capture the complete species gene repertoire. Second, the number of new ‘strain-specific’ genes did not converge to zero upon sequencing of additional strains, at n*=*260, an average of 27 new genes were contributed to the gene pool. Finally, according to Heap’s Law, α values of ≤ 1 are representative of open pangenome. Rarefaction analysis of our pangenome curve using a power-law regression model based on Heap’s Law^28^ showed the pangenome was predicted to be open (*Bpan* (≈ α^28^) = 0.47, curve fit, r^2^=0.999). **(E)** Presence absence variation (PAV) matrix for 260 *C. difficile* genomes is shown alongside a maximum-likelihood phylogeny built from a recombination-adjusted alignment of core genes from Panaroo (2,232 genes, 2,606,142 sites).

Pan-GWAS analysis with Scoary revealed 142 genes with significant clade specificity. Based on KEGG orthology, these genes were classified into four functional categories: environmental information processing (7), genetic information processing (39), metabolism (43), and signalling and cellular processes (53). We identified several uniquely present, absent or organised gene clusters associated with ethanolamine catabolism (C-III), heavy metal uptake (C-III), polyamine biosynthesis (C-III), fructosamine utilisation (C-I, C-III), zinc transport (C-II, C5) and folate metabolism (C-I, C5). A summary of the composition and function of these major lineage-specific gene clusters is given in **Table 2**, and a comparative analysis of their respective genetic architecture can be found in the **Supplementary Data**.

**Table 2.**
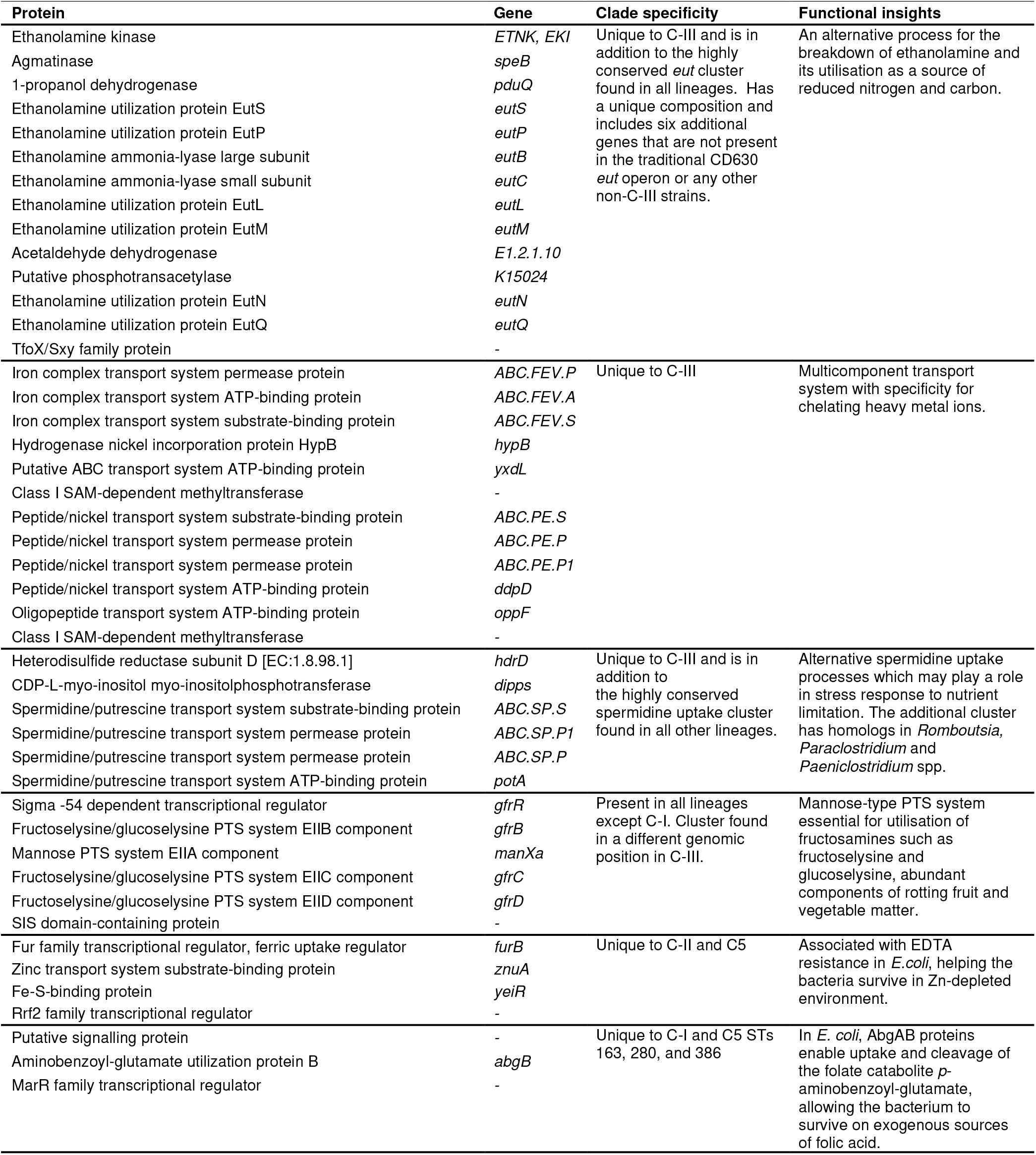
Major clade-specific gene clusters identified by pan-GWAS.

### Cryptic clades CI-III possessed highly divergent toxin gene architecture

Overall, 68.8% (179/260) of STs harboured *tcdA* (toxin A) and/or *tcdB* (toxin B), indicating their ability to cause CDI, while 67 STs (25.8%) harboured *cdtA*/*cdtB* (binary toxin). The most common genotype was A^+^B^+^CDT^-^ (113/187; 60.4%), followed by A^+^B^+^CDT^+^ (49/187; 26.2%), A^-^B^+^CDT^+^ (10/187; 5.3%), A^-^B^-^CDT^+^ (8/187; 4.3%) and A^-^B^+^CDT^-^ (7/187; 3.7%). Toxin gene content varied across clades (C1, 116/149, 77.9%; C2, 35/35, 100.0%; C3, 7/7, 100.0%; C4, 6/34, 17.6%; C5, 18/18, 100.0%; C-I, 2/12, 16.7%; C-II, 1/3, 33.3%; C-III, 2/2, 100.0%) (**Fig. 7**).

**Figure 7.**
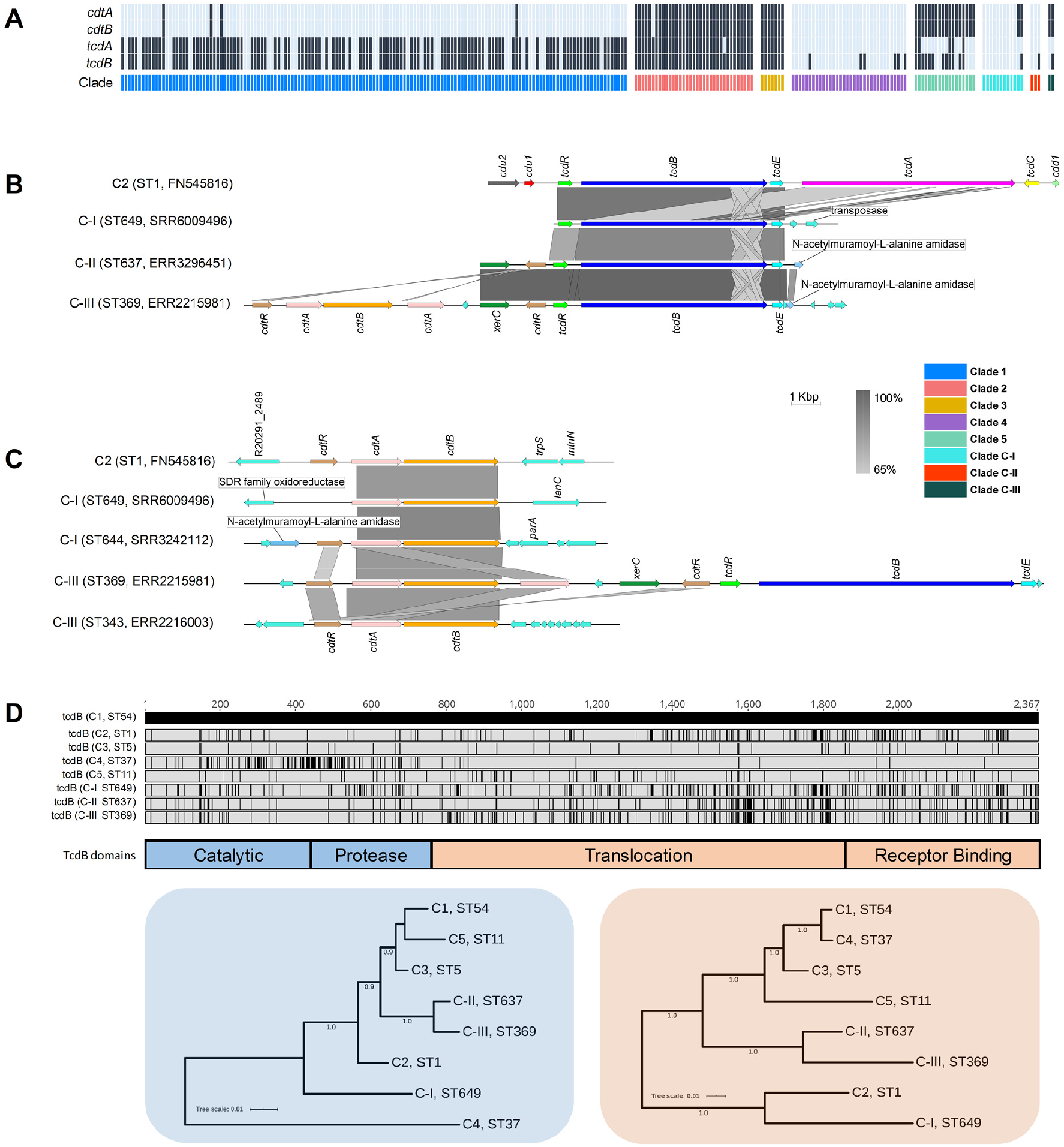
Toxin gene analysis. **(A)** Distribution of toxin genes across *C. difficile* clades (n=260 STs). Presence is indicated by black bars and absence by light blue bars. **(B)** Comparison of PaLoc architecture in the chromosome of strain R20291 (C2, ST1) and cognate chromosomal regions in genomes of cryptic STs 649 (C-I), 637 (C-II), and 369 (C-III). All three cryptic STs show atypical ‘monotoxin’ PaLoc structures, with the presence of syntenic *tcdR, tcdB*, and *tcdE*, and the absence of *tcdA, tcdC, cdd1* and *cdd2*. ST369 genome ERR2215981 shows colocalization of the PaLoc and CdtLoc, see below. **(C)** Comparison of CdtLoc architecture in the chromosome of strain R20291 (C2, ST1) and cognate chromosomal regions in genomes of cryptic STs 649/644 (C-I) and 343/369 (C-III). Several atypical CdtLoc features are observed; *cdtR* is absent in ST649, and an additional copy of *cdtA* is present in ST369, the latter comprising part of a CdtLoc co-located with the PaLoc. **(D)** Amino acid differences in TcdB among cryptic STs 649, 637, and 369 and reference strains from clades C1-5. Variations are shown as black lines relative to CD630 (C1, ST54). Phylogenies constructed from the catalytic and protease domains (in blue) and translocation and receptor-binding domains (in orange) of TcdB for the same eight STs included in (D). Scale bar shows the number of amino acid substitutions per site. Trees are mid-point rooted and supported by 500 bootstrap replicates.

Critically, at least one ST in each of clades C-I, C-II and C-III harboured divergent *tcdB* (89-94% identity to *tcdB*R20291) and/or *cdtAB* alleles (60-71% identity to *cdtA*R20291, 74-81% identity to *cdtB*R20291). These genes were located on atypical and novel PaLoc and CdtLoc structures flanked by mediators of lateral gene transfer (**Fig. 7**). Sequence types 359, 360, 361 and 649 (C-I), 637 (C-II) and 369 (C-III) harboured ‘monotoxin’ PaLocs characterised by the presence of syntenic *tcdR, tcdB* and *tcdE*, and complete absence of *tcdA* and *tcdC*. In STs 360 and 361 (C-I), and 637 (C-II), a gene encoding an endolysin with predicted N-acetylmuramoyl-L-alanine amidase activity (*cwlH*) was found adjacent to the phage-derived holin gene *tcdE*.

Remarkably, a full CdtLoc was found upstream of the PaLoc in ST369 (C-III). This CdtLoc was unusual, characterised by the presence of *cdtB*, two copies of *cdtA*, two copies of *cdtR* and *xerC* encoding a site-specific tyrosine recombinase (**Fig. 7**). Both ST644 (C-I) and ST343 (C-III) were CdtLoc-positive but PaLoc-negative (A^-^B^-^CDT^+^). In ST649 (C-I) *cdtR* was completely absent and, in ST343 (C-III), the entire CdtLoc was contained within the genome of a 56Kbp temperate bacteriophage termed ΦSemix9P1^29^. Toxin regulators TcdR and CdtR are highly conserved across clades C1-5^21^. In contrast, the CdtR of STs 644 (C-I), 343 (C-III) and 369 (C-III) shared only 46-54% amino acid identity (AAI) with CdtR of strain R20291 from clade 2 and ∼40% AAI to each other. Similarly, the TcdR of ST 369 shared only 82.1% AAI compared to R20291 (**Supplementary Data**).

Compared to TcdB of R20291 (TcdBR20291), the shared AAI for TcdBST649_C-I, TcdBST637_C-II and TcdBST369_C-III were 94.0%, 90.5% and 89.4%, respectively. This sequence heterogeneity was confirmed through the detection of five distinct *Hinc*II/*Acc*I digestion profiles of *tcdB* B1 fragments possibly reflecting novel toxinotypes (**Supplementary Data**). TcdB phylogenies identified clade C2 as the most recent common ancestor for TcdBST649_C-I (**Fig. 7**). Phylogenetic subtyping analysis of the TcdB receptor-binding domain (RBD) showed the respective sequences in C-I, C-II and C-III clustered with *tcdB* alleles belonging to virulent C2 strains (**Supplementary Data**). Notably, the TcdB-RBD of ST649 (C-I) shared an AAI of 93.5% with TcdB-RBD allele type 8 belonging to hypervirulent STs 1 (RT027)^13^ and 231 (RT251)^30^. Similarly, the closest match to *tcdB*-RBDs of ST637 (C-II) and ST369 (C-III) was allele type 10 (ST41, RT244)^31^.

## Discussion

Through phylogenomic analysis of the largest and most diverse collection of *C. difficile* genomes to date, we identified major incoherence in *C. difficile* taxonomy, and provide new insight into intra-species diversity and evolution of pathogenicity in this major One Health pathogen.

Our analysis found high nucleotide identity (ANI > 97%) between *C. difficile* clades C1-4, indicating that strains from these four clades (comprising 560 known STs) belong to the same species. This is supported by our core genome and Bayesian analyses, which estimated the most recent common ancestor of *C. difficile* clades C1-4 existed ∼1.61 mya. After this point, there appears to have been rapid population expansion into the four closely related extant clades described today, which include many of the most prevalent strains causing healthcare-associated CDI worldwide^11^. On the other hand, ANI between C5 and C1-4 is on the borderline of the accepted species threshold (95.9-96.2%) and their common ancestor existed 3.89 mya, over 2 Ma before C1-4 diverged. This degree of speciation likely reflects the unique ecology of C5 – a lineage comprising 33 known STs which is well established in non-human animal reservoirs worldwide and recently associated with CDI in the community setting^32^. We identified major taxonomic incoherence among the three cryptic clades and C1-5, evident by ANI values well below the species threshold (∼91%, C-I; ∼94%, C-II; and ∼89%, C-III). Similar ANI value differences were seen between the cryptic clades themselves, indicating they are as divergent from each other as they are individually from C1-5. This extraordinary level of discontinuity is substantiated by our core genome and Bayesian analyses which estimated the common ancestors of clades C-I, C-II and C-III existed 13, 22 and 48 Ma, respectively, at least 9 to 45 Ma before the common ancestor of C1-5. For context, divergence dates for other pathogens range from 10 Ma (*Campylobacter coli* and *C. jejuni*)^33^, 47 Ma (*Burkholderia pseudomallei* and *B. thailandensis*)^34^ and 120 Ma (*Escherichia coli* and *Salmonella enterica*)^35^. Corresponding whole genome ANI values for these species are 86%, 94% and 82%, respectively (**Supplementary Data**).

Comparative ANI analysis of the cryptic clades with >5000 reference genomes across 21 phyla failed to provide a better match than *C. difficile* (89-94% ANI). Similarly, our revised ANI-based taxonomy of the *Peptostreptococcaceae* placed clades C-I, C-II and C-III between *C. difficile* and *C. mangenotii*, the latter sharing ∼77% ANI. The rate of 16S rRNA divergence in bacteria is estimated to be 1–2% per 50 Ma^35^. Contradicting our ANI and core genome data, 16S rRNA sequences were highly conserved across all 8 clades. This indicates that in *C. difficile*, 16S rRNA gene similarity correlates poorly with measures of genomic, phenotypic and ecological diversity, as reported in other taxa such as *Streptomyces, Bacillus* and *Enterobacteriaceae*^36, 37^. Another interesting observation is that C5 and the three cryptic clades had a high proportion (>90%) of MLST alleles that were absent in other clades (**Supplementary Data**) suggesting minimal exchange of essential housekeeping genes between these clades. Whether this reflects divergence or convergence of two species, as seen in *Campylobacter*^38^, is unknown. Taken together, these data strongly support the reclassification of *C. difficile* clades C-I, C-II and C-III as novel independent *Clostridioides* genomospecies. There have been similar genome-based reclassifications in *Bacillus*^39^, *Fusobacterium*^40^ and *Burkholderia*^41^. Also, a recent Consensus Statement^42^ argues that the genomics and big data era necessitate easing of nomenclature rules to accommodate genome-based assignment of species status to nonculturable bacteria and those without ‘type material’, as is the case with these genomospecies.

The NCBI SRA was dominated by C1 and C2 strains, both in number and diversity. This apparent bias reflects the research community’s efforts to sequence the most prominent strains causing CDI in regions with the highest-burden, e.g. ST 1 from humans in Europe and North America. As such, there is a paucity of sequenced strains from diverse environmental sources, animal reservoirs or regions associated with atypical phenotypes. Cultivation bias - a historical tendency to culture, preserve and ultimately sequence *C. difficile* isolates that are concordant with expected phenotypic criteria, comes at the expense of ‘outliers’ or intermediate phenotypes. Members of the cryptic clades fit this criterion. They were first identified in 2012 but have been overlooked due to atypical toxin architecture which may compromise diagnostic assays (discussed below). Our updated MLST phylogeny shows as many as 55 STs across the three cryptic clades (C-I, n=25; C-II, n=9; C-III, n=21) (**Fig. 2**). There remains a further dozen ‘outliers’ which could either fit within these new taxa or be the first typed representative of additional genomospecies. The growing popularity of metagenomic sequencing of animal and environmental microbiomes will certainly identify further diversity within these taxa, including nonculturable strains^43, 44^.

By analysing 260 STs across eight clades, we provide the most comprehensive pangenome analysis of *C. difficile* to date. Importantly, we also show that the choice of algorithm significantly affects pangenome estimation. The *C. difficile* pangenome was determined to be open (i.e. an unlimited gene repertoire) and vast in scale (over 17000 genes), much larger than previous estimates (∼10000 genes) which mainly considered individual clonal lineages^16, 22^. Conversely, comprising just 12.8% of its genetic repertoire (2,232 genes), the core genome of *C. difficile* is remarkably small, consistent with earlier WGS and microarray-based studies describing ultralow genome conservation in *C. difficile*^11, 45^. Considering only C1-5, the pangenome reduced in size by 12% (2,082 genes); another 519 genes were lost when considering only C1-4. These findings are consistent with our taxonomic data, suggesting the cryptic clades, and to a lesser extent C5, contribute a significant proportion of evolutionarily divergent and unique loci to the gene pool. A large open pangenome and small core genome are synonymous with a sympatric lifestyle, characterised by cohabitation with, and extensive gene transfer between, diverse communities of prokarya and archaea^46^. Indeed, *C. difficile* shows a highly mosaic genome comprising many phages, plasmids and integrative and conjugative elements^11^, and has adapted to survival in multiple niches including the mammalian gastrointestinal tract, water, soil and compost, and invertebrates^32^.

Through a robust Pan-GWAS approach we identified loci that are enriched or unique in the genomospecies. C-I strains were associated with the presence of transporter AbgB and absence of a mannose-type phosphotransferase (PTS) system. In *E. coli*, AbgAB proteins allow it to survive on exogenous sources of folate^47^. In many enteric species, the mannose-type PTS system is essential for catabolism of fructosamines such as glucoselysine and fructoselysine, abundant components of rotting fruit and vegetable matter^48^. C-II strains contained Zn transporter loci *znuA* and *yeiR*, in addition to Zn transporter ZupT which is highly conserved across all eight *C. difficile* clades.

*S. enterica* and *E. coli* harbour both *znuA*/*yeiR* and ZupT loci, enabling survival in Zn-depleted environments^49^. C-III strains were associated with major gene clusters encoding systems for ethanolamine catabolism, heavy metal transport and spermidine uptake. The C-III *eut* gene cluster encoded six additional kinases, transporters and transcription regulators absent from the highly conserved *eut* operon found in other clades. Ethanolamine is a valuable source of carbon and/or nitrogen for many bacteria, and *eut* gene mutations (in C1/C2) impact toxin production *in vivo*^50^. The C-III metal transport gene cluster encoded a chelator of heavy metal ions and a multi-component transport system with specificity for iron, nickel and glutathione. The conserved spermidine operon found in all *C. difficile* clades is thought to play an important role in various stress responses including during iron limitation^51^. The additional, divergent spermidine transporters found in C-III were similar to regions in closely related genera *Romboutsia* and *Paeniclostridium* (data not shown). Together, these data provide preliminary insights into the biology and ecology of the genomospecies. Most differential loci identified were responsible for extra or alternate metabolic processes, some not previously reported in *C. difficile*. It is therefore tempting to speculate that the evolution of alternate biosynthesis pathways in these species reflects distinct ancestries and metabolic responses to evolving within markedly different ecological niches.

This work demonstrates the presence of toxin genes on PaLoc and CdtLoc structures in all three genomospecies, confirming their clinical relevance. Monotoxin PaLocs were characterised by the presence of *tcdR, tcdB* and *tcdE*, the absence of *tcdA* and *tcdC*, and flanking by transposases and recombinases which mediate LGT^20, 21, 21^. These findings support the notion that the classical bi-toxin PaLoc common to clades C1-5 was derived by multiple independent acquisitions and stable fusion of monotoxin PaLocs from ancestral Clostridia^52^. Moreover, the presence of syntenic PaLoc and CdtLoc (in ST369, C-I), the latter featuring two copies of *cdtA* and *cdtR*, and a recombinase (*xerC*), further support this PaLoc fusion hypothesis^52^.

Bacteriophage holin and endolysin enzymes coordinate host cell lysis, phage release and toxin secretion^53^. Monotoxin PaLocs comprising phage-derived holin (*tcdE*) and endolysin (*cwlH*) genes were first described in C-I strains^52^. We have expanded this previous knowledge by demonstrating that syntenic *tcdE* and *cwlH* are present within monotoxin PaLocs across all three genomospecies. Moreover, since some strains contained *cwlH* but lacked toxin genes, this gene seems to be implicated in toxin acquisition. These data, along with the detection of a complete and functional^29^ CdtLoc contained within ΦSemix9P1 in ST343 (C-III), further substantiate the role of phages in the evolution of toxin loci in *C. difficile* and related Clostridia ^53^.

The CdtR and TcdR sequences of the new genomospecies are unique and further work is needed to determine if these regulators display different mechanisms or efficiencies of toxin expression^12^. The presence of dual copies of CdtR in ST369 (C-I) is intriguing, as analogous duplications in PaLoc regulators have not been documented. One of these CdtR had a mutation at a key phosphorylation site (Asp61→Asn61) and possibly shows either reduced wild-type activity or non-functionality, as seen in ST11^54^. This might explain the presence of a second CdtR copy.

TcdB alone can induce host innate immune and inflammatory responses leading to intestinal and systemic organ damage^55^. Our phylogenetic analysis shows TcdB sequences from the three genomospecies are related to TcdB in Clade 2 members, specifically ST1 and ST41, both virulent lineages associated with international CDI outbreaks^13, 31^, and causing classical or variant (*C. sordellii*-like) cytopathic effects, respectively^56^. It would be relevant to explore whether the divergent PaLoc and CdtLoc regions confer differences in biological activity, as these may present challenges for the development of effective broad-spectrum diagnostic assays, and vaccines. We have previously demonstrated that common laboratory diagnostic assays may be challenged by changes in the PaLoc of C-I strains^21^. The same might be true for monoclonal antibody-based treatments for CDI such as bezlotoxumab, known to have distinct neutralizing activities against different TcdB subtypes^57^.

Our findings highlight major incongruence in *C. difficile* taxonomy, identify differential patterns of diversity among major clades and advance understanding of the evolution of the PaLoc and CdtLoc. While our analysis is limited solely to the genomic differences between *C. difficile* clades, our data provide a robust genetic foundation for future studies to focus on the phenotypic, ecological and epidemiological features of these interesting groups of strains, including defining the biological consequences of clade-specific genes and pathogenic differences *in vitro* and *in vivo*. Finally, our findings reinforce that the epidemiology of this important One Health pathogen is not fully understood. Enhanced surveillance of CDI and WGS of new and emerging strains to better inform the design of diagnostic tests and vaccines are key steps in combating the ongoing threat posed by *C. difficile*.

## Methods

### Genome collection

We retrieved the entire collection of *C. difficile* genomes (taxid ID 1496) held at the NCBI Sequence Read Archive [https://www.ncbi.nlm.nih.gov/sra/]. The raw dataset (as of 1^st^ January 2020), comprised 12,621 genomes. After filtering for redundancy and Illumina paired-end data (all platforms and read lengths), 12,304 genomes (97.5%) were available for analysis.

### Multi-locus sequence typing

Sequence reads were interrogated for multi-locus sequence type (ST) using SRST2 v0.1.8^58^. New alleles, STs and clade assignments were verified by submission of assembled contigs to PubMLST [https://pubmlst.org/cdifficile/]. A species-wide phylogeny was generated from 659 ST alleles sourced from PubMLST (dated 01-Jan-2020). Alleles were concatenated in frame and aligned with MAFFT v7.304. A final neighbour-joining tree was generated in MEGA v10^59^ and annotated using iToL v4 [https://itol.embl.de/].

### Genome assembly and quality control

Genomes were assembled, annotated and evaluated using a pipeline comprising TrimGalore v0.6.5, SPAdes v3.6.043, Prokka v1.14.5, and QUAST v2.344^16^. Next, Kraken2 v2.0.8-beta^60^ was used to screen for contamination and assign taxonomic labels to reads and draft assemblies.

### Taxonomic analyses

Species-wide genetic similarity was determined by computation of whole-genome ANI for 260 STs. Both alignment-free and conventional alignment-based ANI approaches were taken, implemented in FastANI^5^ v1.3 and the Python module pyani^61^ v0.2.9, respectively. FastANI calculates ANI using a unique *k*-mer based alignment-free sequence mapping engine, whilst pyani utilises two different classical alignment ANI algorithms based on BLAST+ (ANIb) and MUMmer (ANIm). A 96% ANI cut-off was used to define species boundaries^4^. For taxonomic placement, ANI was determined for divergent *C. difficile* genomes against two datasets comprising (i) members of the *Peptostreptococcaceae* (n=25)^23^, and (ii) the complete NCBI RefSeq database (n=5895 genomes, https://www.ncbi.nlm.nih.gov/refseq/, accessed 14^th^ Jan 2020). Finally, comparative identity analysis of consensus 16S rRNA sequences for *C. mangenotii* type strain DSM1289T^23^ (accession FR733662.1) and representatives of each *C. difficile* clade was performed using Clustal Omega https://www.ebi.ac.uk/Tools/msa/clustalo/.

### Estimates of clade and species divergence

BactDating v1.0.1^62^ was applied to the recombination-corrected phylogeny produced by Gubbins (471,708 core-genome sites) with Markov chain Monte Carlo (MCMC) chains of 10^7^ iterations sampled every 10^4^ iterations with a 50% burn-in. A strict clock model was used with a rate of 2.5×10^−9^ to 1.5×10^−8^ substitutions per site per year, as previously defined by He *et al*.^16^ and Kumar *et al*.^27^. The effective sample sizes (ESS) were >200 for all estimated parameters, and traces were inspected manually to ensure convergence. To provide an independent estimate from BactDating, BEAST v1.10.4^63^ was run on a recombination-filtered gap-free alignment of 10,466 sites with MCMC chains of 5×10^8^ iterations, with a 9×10^−7^ burn-in, that were sampled every 10^4^ iterations. The strict clock model described above was used in combination with the discrete GTR gamma model of heterogeneity among sites and skyline population model. MCMC convergence was verified with Tracer v1.7.1 and ESS for all estimated parameters were >150. For ease of comparison, clade dating from both approaches were transposed onto a single MLST phylogeny. Tree files are available as **Supplementary Data** at http://doi.org/10.6084/m9.figshare.12471461.

### Pangenome analysis

The 260 ST dataset was used for pangenome analysis with Panaroo v1.1.0^64^ and Roary v3.6.0^65^. Panaroo was run with default thresholds for core assignment (98%) and blastP identity (95%). Roary was run with a default threshold for core assignment (99%) and two different thresholds for BlastP identity (95%, 90%). Sequence alignment of the final set of core genes (Panaroo; n=2,232 genes, 2,606,142 bp) was performed using MAFFT v7.304 and recombinative sites were filtered using Gubbins v7.304^66^. A recombinant adjusted alignment of 471,708 polymorphic sites was used to create a core genome phylogeny with RAxML v8.2.12 (GTR gamma model of among-site rate-heterogeneity), which was visualised alongside pangenome data in Phandango^67^. Pangenome dynamics were investigated with PanGP v1.0.1^16^.

Scoary^68^ v1.6.16 was used to identify genetic loci that were statistically associated with each clade via a Pangenome-Wide Association Study (pan-GWAS). The Panaroo-derived pangenome (n=17,470) was used as input for Scoary with the evolutionary clade of each genome depicted as a discrete binary trait. Scoary was run with 1,000 permutation replicates and genes were reported as significantly associated with a trait if they attained *p-*values (empirical, naïve and Benjamini-Hochberg-corrected) of ≤0.05, a sensitivity and specificity of > 99% and 97.5%, respectively, and were not annotated as “hypothetical proteins”. All significantly associated genes were reannotated using prokka and BlastP and functional classification (KEGG orthology) was performed using the Koala suite of web-based annotation tools^69^.

### Comparative analysis of toxin gene architecture

The 260 ST genome dataset was screened for the presence of *tcdA, tcdB, cdtA* and *cdtB* using the Virulence Factors Database (VFDB) compiled within ABRicate v1.0 [https://github.com/tseemann/abricate]. Results were corroborated by screening raw reads against the VFDB using SRST2 v0.1.8^58^. Both approaches employed minimum coverage and identity thresholds of 90 and 75%, respectively. Comparative analysis of PaLoc and CdtLoc architecture was performed by mapping of reads with Bowtie2 v.2.4.1 to cognate regions in reference strain R20291 (ST1, FN545816). All PaLoc and CdtLoc loci investigated showed sufficient coverage for accurate annotation and structural inference. Genome comparisons were visualized using ACT and figures prepared with Easyfig^21^. MUSCLE-aligned TcdB sequences were visualized in Geneious v2020.1.2 and used to create trees in iToL v4.

### Statistical analyses

All statistical analyses were performed using SPSS v26.0 (IBM, NY, USA). For pangenome analyses, Chi-squared test with Yate’s correction was used to compare the proportion of core genes and a One-tailed Mann-Whitney U test was used to demonstrate the reduction of gene content per genome, with a p-value ≤ 0.05 considered statistically significant.

## Author contributions

D.R.K., K.I., D.W.E., and T.V.R. designed the study. D.R.K., K.I., C.R., B.K., E.G.A., and K.E.D. performed experimental work. D.R.K., K.I., C.R., B.K., E.G.A., D.P.S., X.D., K.E.D., D.W.E., C.R., and T.V.R. analysed data and drafted the manuscript. All authors edited and approved the final version of the manuscript. The corresponding author had full access to all the data in the study and had final responsibility for the decision to submit for publication.

## Acknowledgements

This work was supported, in part, by funding from The Raine Medical Research Foundation (RPG002-19) and a Fellowship from the National Health and Medical Research Council (APP1138257) awarded to D.R.K. K.I. is a recipient of the Mahidol Scholarship from Mahidol University, Thailand. This work was also supported by EULac project ‘Genomic Epidemiology of *Clostridium difficile* in Latin America (T020076)” and by the Millennium Science Initiative of the Ministry of Economy, Development and Tourism of Chile, grant ‘Nucleus in the Biology of Intestinal Microbiota” to D.P.S. This research used the facilities and services of the Pawsey Supercomputing Centre [Perth, Western Australia] and the Australian Genome Research Facility [Melbourne, Victoria].

## Additional information

Supplementary Data is available at http://doi.org/10.6084/m9.figshare.12471461

